# TCR repertoire analysis quantifies effect of irradiation to homeostasis of iNKT cell development in the thymus of mice

**DOI:** 10.1101/2025.05.12.653114

**Authors:** Kazumasa B. Kaneko, Mio Hayama, Nanako Uchida, Takahisa Miyao, Nobuko Akiyama, Taishin Akiyama, Tetsuya J. Kobayashi

## Abstract

Thymic T cell development shows homeostasis in which T cell production recovers after temporal impairment by various stressors. Invariant natural killer T (iNKT) cells in the thymus contribute to recovery of thymic development after irradiation owing to their greater tolerance to irradiation compared to conventional thymocytes. However, whether and how iNKT cell development recovers from irradiation remains unknown. Here we show that iNKT cells in the thymus exhibits much slower post-irradiation recovery than conventional thymocytes. We compared T cell receptor (TCR) alpha chain repertoire of the thymus in 42 days after irradiation and quantified the sustained decline in the number of iNKT cells against other thymocytes. We found that fluctuation of V and J genes usage does not correlate with the number iNKT cells suggesting that differentiation process of iNKT cells after TCR gene rearrangement is impaired by irradiation. Mathematical modelling of recovery dynamics of iNKT cells implied that lack of rapid proliferation immediately after irradiation unlike conventional thymocytes contributes to the prolonged reduction of iNKT cell population. These findings may suggest possibility on association between the prolonged reduction of iNKT cells after irradiation and autoimmune diseases caused by irradiation in bone marrow transplantation.

## 1 Introduction

The thymus is the primary organ responsible for producing appropriate repertoires of T cells (Takahama 2006). Even with its role for immuno-homeostasis, the thymus is relatively susceptible to various stressors, including mental stresses, viral infections, irradiation, and other stimuli (Boehm and Swann 2013; Gruver and Sempowski 2008). While the thymus of a healthy individual is capable of recuperating promptly from such insults, prolonged recovery interval would lead to diminished production of T-cells during this interval, thereby increasing the likelihood of compromised T-cell-mediated immune responses (Gruver and Sempowski 2008; Parkman and Weinberg 1997).

The mechanism of thymic homeostasis to disturbances has been investigated utilizing a variety of measurement techniques. In previous studies using flow cytometric analysis, recoveries of both conventional thymocytes and thymic epithelial cells (TECs) have been focused on because their cross-talk is critical for the thymic development (Abramson and Anderson 2017). The integration of statistical inference and mathematical modeling with these quantitative datasets has also elucidated the dynamic aspects of the homeostatic recovery of the damaged thymus through the interaction between thymocytes and TECs (Robert et al. 2021; Kaneko et al. 2019).

The differentiation mechanisms of various unconventional T cell subsets in the thymus differs markedly from that of conventional T cells, potentially leading to distinct homeostatic mechanisms and dynamics during recovery from the damages (Pellicci, Koay, and Berzins 2020). However, the extent to which unconventional thymic T cell subpopulations respond to disturbances and whether they exhibit the same degree of homeostatic resilience as conventional T cells has not yet been fully investigated.

Among the subpopulations, invariant natural killer T cells (iNKT cells) and their homeostatic production in the thymus warrant particular attention partially because of their potential relations with obesity, asthma, and tumor immunity (Satoh and Iwabuchi 2018; Yip et al. 2019). The Inkt cells are distinguished from the conventional T cells by their specific TCRα gene combination of the TRAV11 (Vα14) and TRAJ18 (Jα18), resulting in the invariant complementarity-determining region 3 (CDR3) sequence (Greenaway et al. 2013). Moreover, while the selection of conventional thymocytes depends on interactions with self-antigen peptides loaded onto major histocompatibility complex (MHC) molecules presented by thymic professional antigen-presenting cells such as TECs, the differentiation and selection process of iNKT cells is distinct. The fate determination of iNKT cells requires interaction with endogenous lipid antigens presented by CD4+CD8+ double-positive (DP) thymocytes through CD1d molecules (Pellicci, Koay, and Berzins 2020).

Recent studies, including those by Cosway et al. have indicated that iNKT cells demonstrate a heightened tolerance to irradiation compared to other thymocyte populations, a factor potentially responsible for the recovery of T cells immediately after irradiation (Cosway et al. 2022). Cosway et al. also observed that, in iNKT cell-deficient mice, the reduced population sizes of thymocyte subtypes by irradiation stay low after irradiation compared to the wild-type (WT) counterparts. Nonetheless, it remains unclear whether the sizes of individual T-cell subtypes in the WT thymus, especially that of iNKT cells, actually revert to pre-irradiation levels or not. Moreover, the impacts of irradiation on the iNKT cell-specific TCR gene rearrangement, along with their subsequent maturation processes involving antigen presentation, remain largely unexplored.

To address these problems, we quantified the temporal changes in iNKT cell counts within the thymus of WT mice during the recovery process from irradiation, utilizing Fluorescence-Activated Cell Sorting (FACS). We observed that thymic iNKT cell counts did not recover for 40 days after irradiation. Additionally, we analyzed the repertoire of TCRα chains in thymocytes after irradiation. Prior to irradiation, the iNKT cell-specific sequences occupies a predominant fraction in both the DP and CD4+ single positive (SP4) subsets. Although this proportion briefly surged immediately after irradiation, it later declined and failed to return to pre-irradiation levels, confirming the trends observed in FACS analyses. Notably, this reduction in the proportion of iNKT-specific sequences was more pronounced than the alterations observed in other representative sequences in response to irradiation.

While we observed that the genomic distance between selected V and J genes after irradiation fluctuated around its pre-irradiation level in correlation with the minor variations in iNKT cell population size, it did not explain the failure of iNKT cell numbers to return to their original levels.

In exploring the underlying cause for the non-recovery of the thymic iNKT cell population, we examined the DP cell proliferation, the major mechanism of thymocyte recovery after irradiation (Kaneko et al. 2019). We constructed a mathematical model by assuming that DP iNKT cell precursors temporarily proliferate after irradiation as normal DP cells do and found that the predicted dynamics from the model could not reproduce the observed population dynamics of thymocytes.

This result suggests that DP iNKT cell precursors may lack the ability for transient upregulation of proliferation following irradiation, unlike what has been verified for normal DP thymocytes.

## 2 Results

### 2.1 iNKT cell counts declined and did not recover after irradiation

We aim to investigate the long-term effects of acute sub-lethal total body irradiation on iNKT cell populations in the thymus of mice. In this study, 7-week-old female Balb/cA mice were exposed to a sub-lethal irradiation dose of 4.5 Gy. Under these conditions, the thymus undergoes transient involution, with maximum shrinkage observed around 4 days post-irradiation (Kaneko et al. 2019). We assessed the long-term effects of sub-lethal dose irradiation at three time points: day 13, day 30, and day 42, in addition to pre-irradiation. As expected, thymic cell numbers were restored by day 13 and reduced again by day 30, likely due to the reported bimodal effect (Takada et al. 1969; Xiao et al. 2017) compared to age-matched controls (Supplementary Figure 1). By day 42, total thymocyte numbers were restored to near control levels.

In the flow cytometric analysis, we initially characterized the post-irradiation dynamics of CD4-CD8-double negative (DN), DP, and SP4 thymocyte fractions (Figure 1), which encompass the majority of iNKT cells and their precursors. Data analysis showed a substantial recovery in the numbers of SP4 and DP cells by day 13 post-irradiation (Figure 2A) After a subsequent reduction in these populations on day 30, a secondary phase of recovery was observed by day 42 (Figure 2A). This pattern aligns with the bimodal effect of sub-lethal irradiation on thymocytes. In contrast, the number of DN cells was not recovered on day 13 but exhibited some recovery thereafter (Figure 2A).

**Figure 1.**
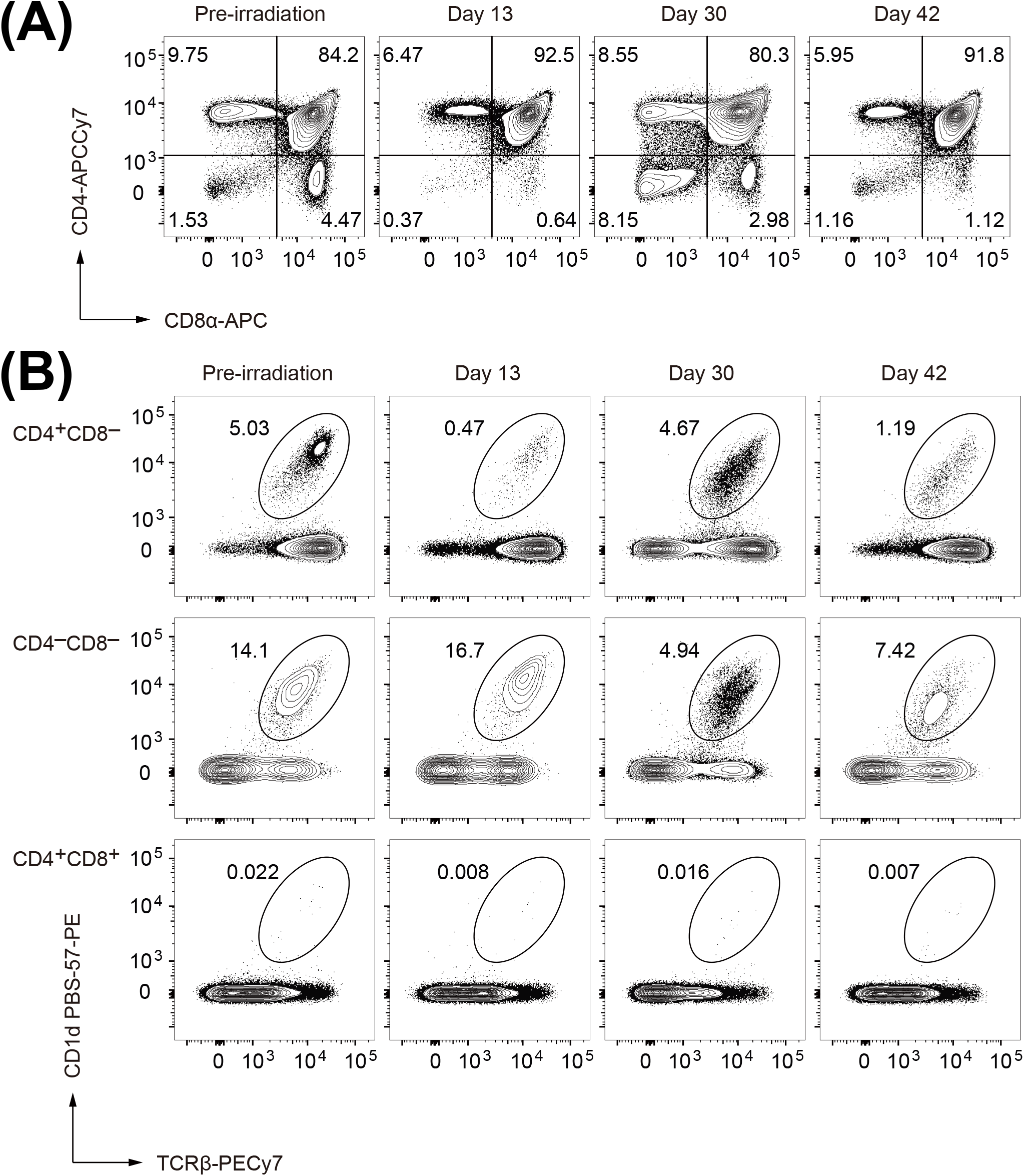
Typical flow cytometric profiles of the thymocytes after the sub-lethal dose radiation. Percentatge of each fraction is shown in the panel. (A) Thymocytes were analyzed by staining with anti-CD4 and anti-CD8α. (B) iNKT cells in each thymocyte subset are defined as CD1d PBS-57-PE+ cells.

**Figure 2.**
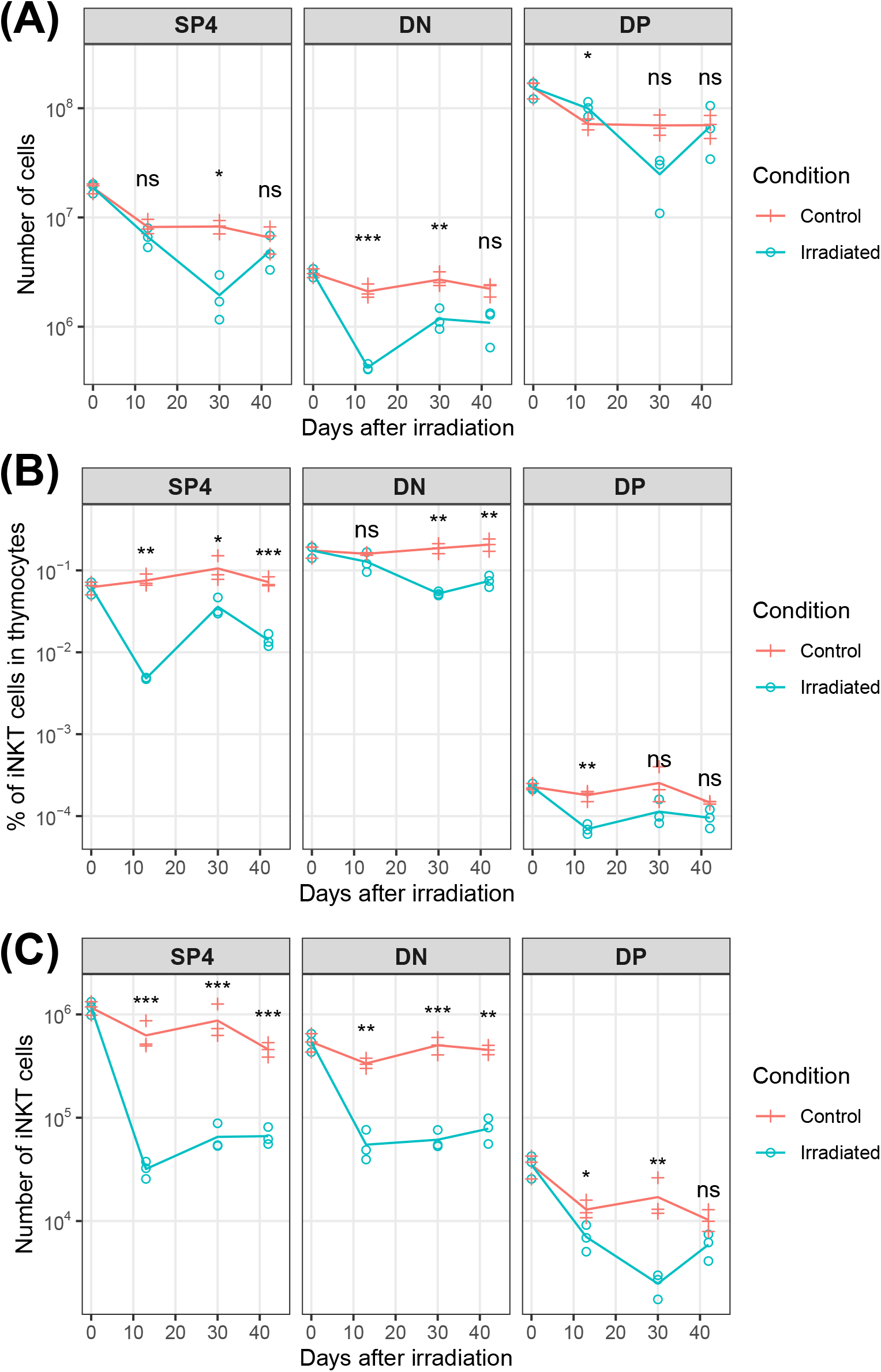
Kinetics of thymocyte subsets after sub-lethal irradiation. Biological triplicate samples were collected and analyzed at each time point before and after irradiation and (A) the number of SP4, DN, and DP thymocytes, (B) proportions of iNKT cells to thymocytes in SP4, DN, and DP subsets, and (C) the number of SP4, DN, and DP iNKT cells were counted. Unequal variance t-test was conducted between irradiated and control groups at each time point. ns: p ≥ 0.05, *: p < 0.05, **: p < 0.01, ***: p < 0.001.

Investigating the impact of irradiation on thymic iNKT cell dynamics, we utilized staining with PBS-57 ligand complexed to CD1d tetramers (Liu et al. 2006), in conjunction with TCRβ antibody, to assess iNKT cell frequencies in these thymocyte subsets. Analysis revealed notable fluctuations in iNKT cell percentages post-irradiation. Specifically, percentages of iNKT cells in the SP4 and DP subsets exhibited reductions on day 13, followed by an increase on day 30, succeeded by another decline (Figure 2B). Notably, accurate enumeration of DP iNKT cell precursors via flow cytometric analysis may have been compromised due to their markedly low frequency within DP thymocytes.

Conversely, the percentage of iNKT cells in the DN subset remained consistent with pre-irradiation levels on day 13 but reduced thereafter (Figure 2B). Notably, analysis of iNKT cell numbers within each subset (Figure 2C) revealed reductions in both SP4 and DN subsets by day 13, persisting until day 42. In the DP subset, the cell number of iNKT precursor cells decreased by day 30, with a modest recovery observed by day 42. However, it is pertinent to note the limited abundance of iNKT cells in the DP subset.

Our flow cytometric analysis unveiled a distinct recovery pattern for thymic iNKT following acute sublethal total body irradiation, contrasting conventional thymocytes. Notably, iNKT cell numbers within the SP4 and DN thymocyte fractions exhibited a persistent decrease for an extended period post-irradiation. This unique recovery kinetics would create a disproportionate reduction and recovery in the ratios of iNKT cells in the SP4 and DN fractions after the radiation.

### 2.2 The iNKT cells were greatly reduced in the TCRα chain repertoire after irradiation

To further investigate dynamics of the iNKT populations, we utilized TCR repertoire measurement. Contrary to the diverse TCR repertoires observed in normal T cells, the TCRα chain of iNKT cells is invariant, whose V and J genes are Vα14 (TRAV11/TRAV11D) and Jα18 (TRAJ18), respectively.

Moreover, the CDR3 region is defined by the sequence CVVGDRGSALGRLHF, which we designate as the iNKT cell sequence. Exploiting this specificity, we can quantify and trace the temporal variation of iNKT cells by sequencing.

The observed variation in the proportion of iNKT cell sequences within the total sequence counts post-irradiation was consistent with that identified through FACS under identical experimental conditions (as depicted in Figure 3A). Notably, there was a significant increase in the percentage of iNKT cells at day 7 after irradiation, which was not measure by FACS. This increase aligns with the observations of Cosway et al., reporting an elevation in the percentage of iNKT cells at day 1 following irradiation (Cosway et al. 2022). Additionally, the proportions of iNKT cells within each cell type and the distribution of each cell type within the iNKT cell population were consistent with previous studies summarized in Table 1. These results collectively suggest that the proportion of the iNKT cell sequence in the TCRα repertoire tends to be proportional to the fraction of iNKT cells in thymocytes.

**Figure 3.**
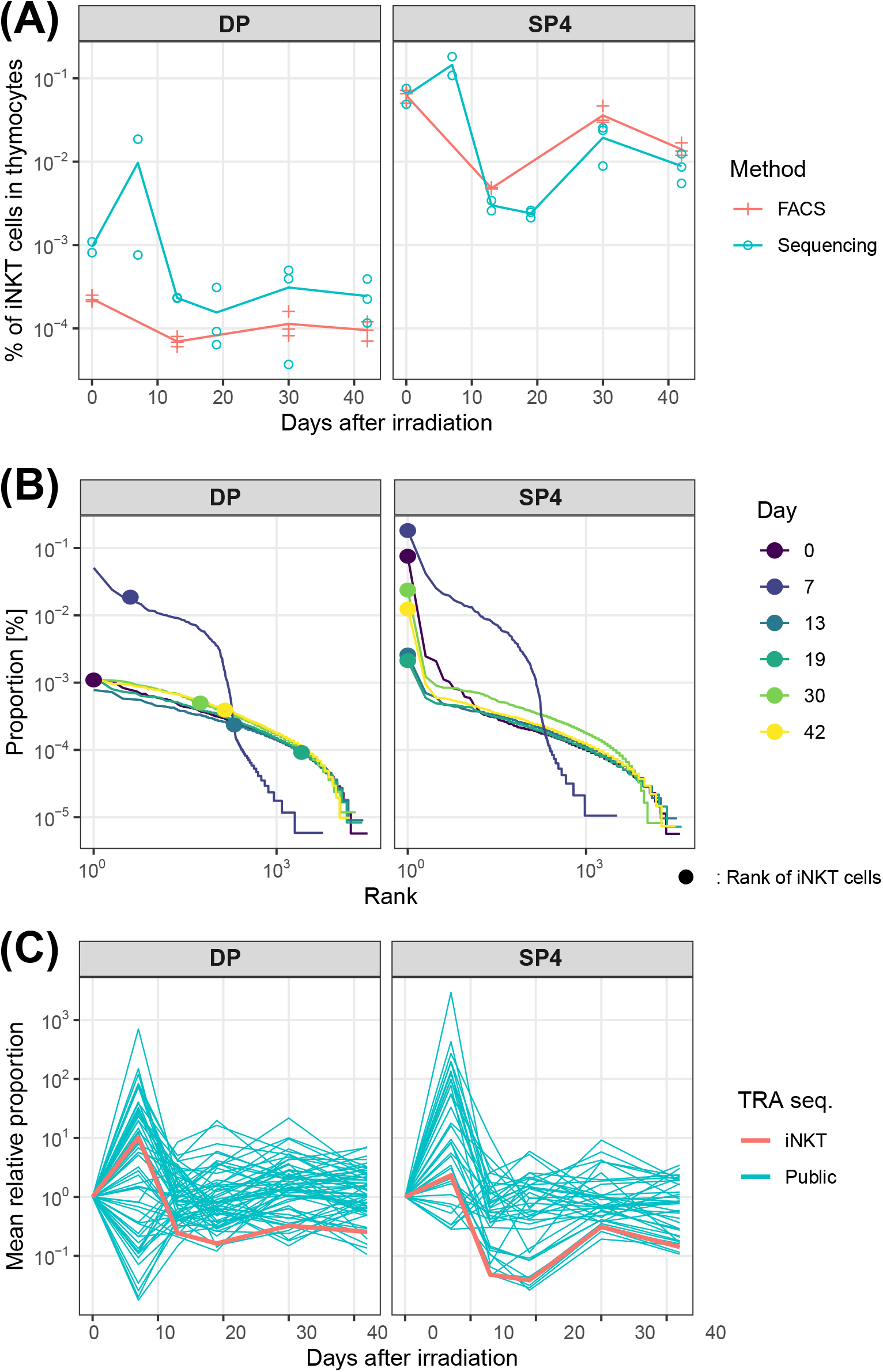
Kinetics of TCRα repertoire of thymocytes after sub-lethal irradiation. (A) Comparison between FACS and repertoire sequencing on kinetics of proportion of iNKT precursors to thymocytes in SP4 and DP subsets. For repertoire sequencing, biological duplicate samples were collected and analyzed were analyzed on 0, 7, and 13 days and triplicate were analyzed on 19, 30, and 42 days after irradiation. (B) Rank-size distribution of TCRα sequence counts of thymocytes. Cirlces represent position of iNKT cells. (C) Kinetics of mean ratio of public sequences and the iNKT cell sequence to the day before irradiation. Public sequences are defined as the sequences that were observed in all samples before irradiation and at least 1 sample at each day after irradiation.

**Table 1.** Comparison of proportion of iNKT cells with those from previous studies.

|  | Strain | Method | iNKT in<br>thymocytes<br>[%] | iNKT<br>in<br>SP4<br>[%] | iNKT<br>in<br>DP<br>[%] | SP4<br>in<br>iNKT<br>[%] | DN<br>in<br>iNKT<br>[%] | DP<br>in<br>iNKT<br>[%] |
| --- | --- | --- | --- | --- | --- | --- | --- | --- |
| Our result | Balb/cA | Sequencing | N/A | 6.2 | 0.1 | N/A | N/A | N/A |
| Our result | Balb/cA | FACS | 0.97 | 6.2 | 0.02 | 67 | 31 | 2 |
| (White et al. 2014) | C57BL/6 | FACS | 0.32 | N/A | N/A | N/A | N/A | N/A |
| (Gapin et al. 2001) | C57BL/6 | FACS | 0.7 | N/A | N/A | 50 | 27 | 0.1 |
| (Lee et al. 2009) | C57BL/6 | FACS | 0.4 | N/A | N/A | 65 | 31 | 3.6 |

Then, we delved into the alterations in the TCRα repertoire following irradiation, as detailed in Figure 3B. Prior to irradiation, the iNKT cell sequence was represented as the most predominant and abundant sequence within the SP4 repertoire and also ranked high among the DP repertoires. Post-irradiation, while the iNKT sequence continued to be the most abundant in SP4 repertoire, its fraction in SP4 repertoire first declined and then partially recovered compared to the pre-irradiation level, but not fully restored. In contrast, iNKT sequence in DP repertoire dropped in ranking and did not recover at all. The result for SP4 iNKT cells is consistent with the observation by FACS in Figure 2B. Given the high sensitivity of sequencing, the result for DP iNKT cells is expected to represent actual dynamics, which was not captured by FACS.

While the ranking of iNKT cells did not recover, Figure 3B shows that the overall repertoire immediately recuperates to its original pre-irradiation distribution by day 13 from the significant distortion observed at day 7 post-irradiation. Thus, X-ray irradiation transiently influences the diversity of the TCRα repertoire, but its effects are not enduring. To scrutinize the impact of irradiation on thymocytes possessing distinct TCRs, we conducted a comparative analysis of the temporal variation in the frequency of TCR sequences ubiquitously detected across multiple samples. Specifically, we defined the common sequences as those identified in both of the two samples before irradiation and also in at least one sample on each subsequent day after irradiation. The proportion of certain common sequences in the repertoire exhibited an increase, while others displayed a decrease relative to pre-irradiation levels (Figure 3C). This pattern suggests that irradiation exerts a non-specific, stochastic perturbation on the proportions of these common sequences, which may lead to quick recovery of the overall repertoire distribution. In contrast, the iNKT sequences were among those most significantly diminished when compared to the trajectories of common sequences, implying the specific mechanism that prevents the recovery of iNKT cells.

A simple explanation of the iNKT cell-specific imperfect recovery is that irradiation impairs the thymus’s capacity to produce iNKT cells. Given the pairing of TRAV11 and TRAJ18 in the iNKT cell sequence as the selected VJ genes for the second VJ recombination (Hu et al. 2010),we posited that this VJ recombination process might be implicated in the observed diminution of iNKT cell populations. To substantiate this hypothesis, we conducted an analysis of the variability in VJ gene usage, finding that an elevated preference at day 19 post-irradiation for selecting more proximal VJ gene pairs on genome than those at day 0. This trend switched towards a more frequent selection of distally located VJ gene pairs at day 30, followed by a reversion to the selection of proximal pairs at day 42 (Figure 4A). In essence, thymocytes exhibiting TCRα chains resulting from the initial VJ recombination were more abundant at days 19 and 42, whereas those with TCRα chains undergoing multiple VJ recombinations were more prevalent at day 30. This pattern is consistent with our observation that the fraction of the iNKT cell sequence generated by the second VJ reconstruction was higher at day 30 than at day 19 or day 42 after irradiation (Figure 4B). Nevertheless, these observed trends do not sufficiently explain the persistent reduction and lack of recovery in the iNKT cell sequences.

**Figure 4.**
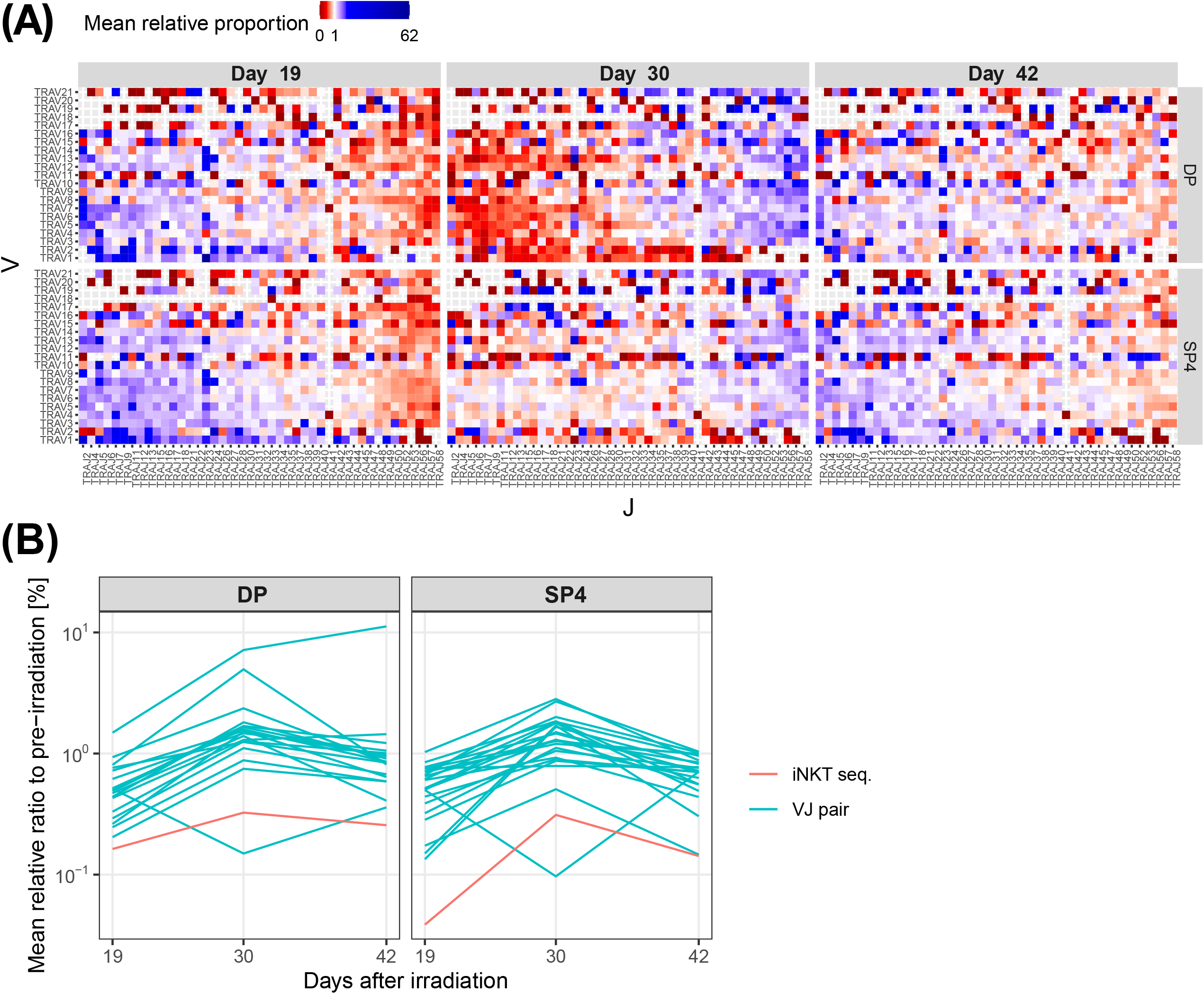
Dynamics of mean relative VJ usages after irradiation when values before irradiation are normalized to 1. (A) Comparison of DP and SP at 19, 30, 42 days after irradiation. (B) Comparison of VJ usages (blue) with the mean relative proportion of the iNKT cell sequence (red). The pairs of top 5 distal VJ genes are extracted.

### 2.3 DP iNKT cell precursors are likely not to transiently proliferate unlike DP thymocytes

Given that the observed alternations in VJ gene usage prompted by irradiation are not sufficient to account for the sustained reduction in iNKT cells, this decline may be associated with the intrinsic differences in differentiation kinetics of iNKT cells from conventional thymocytes. In the experiment in this work, the number of DP thymocytes almost recovered at day 13 after irradiation (Figure 2A). This observation aligns with our previous experiments using the identical experimental system (Kaneko et al. 2019), wherein we elucidated that DP thymocytes exhibit a rapid resurgence, primarily facilitated by transient upregulation of proliferation upon irradiation. Thus, if iNKT cells are deficient in this specific attribute, the sustained reduction can be explained. Nevertheless, considering the exceedingly minute proportion of iNKT precursors within DP thymocytes, even if iNKT DP precursors were capable of a transient proliferative response akin to that of normal DP thymocytes, the possibility remains that such a diminutive fraction may not effectively rebound in the face of competitive recovery dynamics with the predominant majority of other DP thymocytes. We investigate the latter possibility by simplifying and extending our proposed mathematical model (Kaneko et al. 2019) used for reproducing the recovery dynamics of DP and other thymocytes.

Let the number of DP thymocytes be *n*_1_ and the number of DP iNKT cell precursors be *n*_1_, and we describe the population dynamics of their cell numbers by the following ordinary differential equations (ODEs):

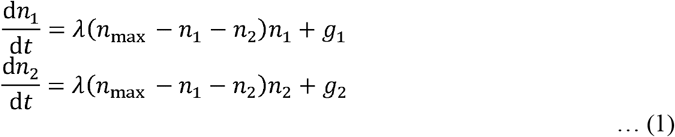

The first term in each equation represents the marginal proliferation, which is negatively regulated by the total number of DP thymocytes as we clarified in our previous work (Kaneko et al. 2019).

Because of this negative regulation, the proliferation is upregulated when the total number of DP cells temporarily decreases far below the steady-state number. We here assume that both DP thymocytes and DP iNKT cell precursors have the same regulatory mechanism for proliferation. The second terms, *g*_1_ and *g*_2_, represent the influx of new cells into DP thymocytes and DP iNKT cell precursors, respectively. The effects of interactions between TECs and other cells are abbreviated here for simplicity.

Figure 5 shows the simulated population dynamics of DP thymocytes and DP iNKT cell precursors by this model. By fixing the initial population size of the DP thymocytes at 10% of the steady-state value, we examined the behavior of iNKT cell precursors starting from different initial population sizes. When the initial iNKT cell count was 10% of its steady-state value, iNKT cell numbers initially overshot and then slowly returned to the the steady state level (Figure 5A). By contrast, when the initial count was reduced to 0.1%, the number of iNKT cells initially returned up to a certain percentage of the steady-state level, then the rate of recovery substantially was slowed down (Fig. 5b). Intuitively, this slowing down happens because the DP cells that recovered faster than iNKT DP cells suppress their proliferation.

**Figure 5.**
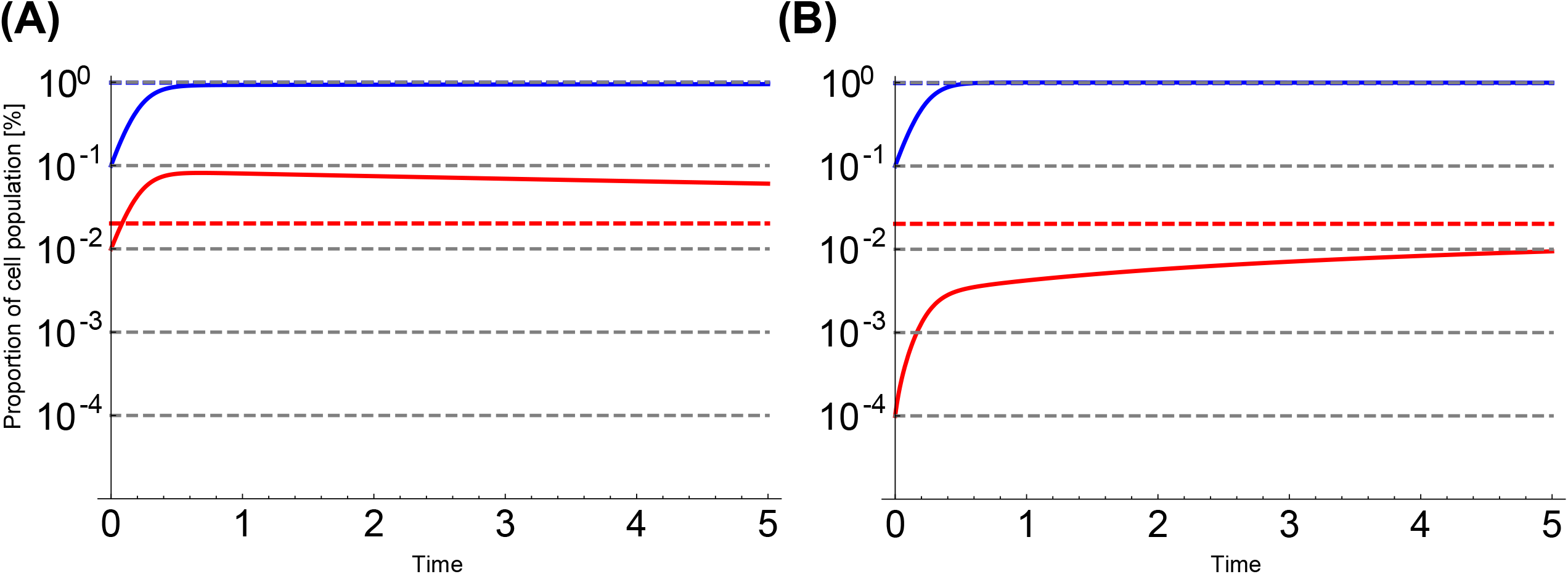
Trajectories of the mathematical model of thymocytes population. DP thymocytes: blue, DP iNKT precursors: red. Trajectories of the mathematical model: solid line, steady states: dotted line. The initial values of DP iNKT precursors are set to (A) 10% and (B) 0.1% of DP thymocytes.

The latter result indicates that the prolonged reduction of DP iNKT precursors can be explained by the recovery competition alone. However, the latter scenario requires that the DP iNKT precursor population diminishes more than the other DP cells immediately irradiation, which is inconsistent with previous studies showing that iNKT cells are more resistant to irradiation (Cosway et al. 2022). Thus, our modeling analysis suggests that DP iNKT cells do not possess the mechanism for transient proliferation or that irradiation may have caused irreversible damage to iNKT cell differentiation, which slowed the recovery of their population size.

## 3 Discussion

In this study, we employed FACS alongside sequencing of TCRα chains to delineate temporal alterations in the population of intra-thymic iNKT cells in mice subjected to X-ray irradiation. We found that irradiation induced a reduction in the intra-thymic iNKT cell count, which persisted for six weeks. Additionally, our sequencing analysis verified that this decrement in intra-thymic iNKT cell counts is not attributable to genetic reconstitution because the patterns of VJ usage variation deviated from the trend in the reduced intra-thymic iNKT cell counts. Using mathematical modeling, we also excluded the possibility that the sustained reduction is due to the proliferation competition of DP iNKT precursors with other DP thymocytes. These results suggest that the prolonged reduction of iNKT cells is attributable to their distinct developmental process from conventional thymocytes and impairments in the process. Finally, we discuss the possible steps and factors in the iNKT developmental process.

The iNKT cells undergo maturation through interactions with a diverse array of cells in the thymus. The iNKT cells require CD1d-mediated antigen presentation by DP thymocytes in the early stages of iNKT cell differentiation (Pellicci, Koay, and Berzins 2020). Additionally, while iNKT cells possess invariant α-chains, their reactivity to self-antigens varies depending on the β-chain. Consequently, only those cells that exhibit self-tolerance can mature through negative selection mediated by antigen presentation from dendritic cells (Bedel et al. 2014). In addition to antigen presentation to TCR, cytokine signaling from mTECs and thymic tuft cells contribute to the development of iNKT cells (White et al. 2014; Lucas et al. 2020). Irradiation might impair these signaling processes, resulting in an increase in iNKT cells undergoing cell death during maturation.

Moreover, iNKT cells demonstrate a markedly elevated proliferation rate during CD44-stage 1 (Föhse et al. 2013; Klibi, Amable, and Benlagha 2020). The dysfunctions in certain intracellular signaling pathways involved in this proliferation might contribute to the observed reduction in iNKT cell populations. The iNKT cells encompasses both immature variants, which are subsequently released into the periphery, and mature subsets that reside long-term within the thymus (Berzins et al. 2006). A potential determinant in the failure of thymic iNKT cell counts in recovery could be the loss of essential factors that retain iNKT cells within the thymus, leading to an elevated release of these cells into peripheral tissues. Investigating the β-chain repertoire of residual iNKT cells in the thymus, alongside analyses of their subtypes and proliferation state, and CD1d expression in surrounding cells, might identify the loci of developmental anomalies in iNKT cells.

To investigate the detailed developmental process of iNKT cells and the impairment caused by irradiation, it is necessary to employ a method capable of accurately collecting and analyzing the small fraction of iNKT-related cells within thymic cells. The search for highly specific iNKT markers and the utilization of high-throughput FACS are essential. Additionally, simultaneous analysis of the repertoire and phenotype via single-cell sequencing will become increasingly important due to its high sensitivity in identifying cells with unique TCR sequences as demonstrated in this paper. Apart from its mechanisms, the long-term reduction in iNKT cell counts induced by irradiation raises a couple of clinical implications. Defects in iNKT cells have been implicated in a diverse array of diseases, encompassing autoimmune disorders (Morris et al. 2009), obesity (Satoh and Iwabuchi 2018), asthma (Yip et al. 2019), and cancer (Vivier et al. 2012). However, the mechanistic underpinnings of these associations remain largely elusive. A contributing factor to this knowledge gap is the reliance on Jα18 knockout mice as a model for iNKT cell deficiency (Zhang et al. 2016; Chandra et al. 2015). These mice exhibit a skewed TCRα chain repertoire, leading to an altered immuno-repertoire landscape that extends beyond iNKT cells. The development of novel methodologies isolating the impact of iNKT cell deficits offers promising avenues for elucidating their role in various diseases. This includes assessing whether the array of peripheral immune alterations attributable to iNKT cell deficiency can be recapitulated in models of irradiation-induced iNKT cell depletion. Such investigations are pivotal in advancing our comprehension of iNKT cell functionality and their disease associations.

Moreover, the employment of irradiation in bone marrow transplantation procedures has been associated with the onset of autoimmune diseases (Alawam et al. 2022). The impact of irradiation on the immune system may be further clarified by examining whether the immune abnormalities are alleviated by transplantation of iNKT cells after irradiation. Such studies are essential for delineating the role of iNKT cells in modulating immune responses and in the potential mitigation of autoimmune diseases following bone marrow transplantation.

In summary, we have shown that X-irradiation in mice models leads to a sustained reduction in iNKT cells within the thymus, persisting for at least six weeks post-irradiation. Future research should focus on identifying the specific junctures in iNKT cell development that are impacted by irradiation. Building upon the results of this study, the causal relationship between the diminution of iNKT cells and disease can be further explored using irradiation models. Additionally, it is imperative to investigate whether the reduction in iNKT cells directly contributes to the onset of diseases associated with irradiation, thereby enhancing our understanding of the immunological consequences of irradiation and its clinical implications for disease development.

## 4 Experimental Procedures

### 4.1 Mice

Animal experiments were performed in accordance with the Guidelines of the Institutional Animal Care and Use Committee of RIKEN, Yokohama Branch (2018-075). Balb/cA Jcl mice (CLEA Japan) were maintained in standard controlled conditions with a 12-h lighting cycle and access to chow and water ad libitum. Seven-week-old female mice received 4.5 Gy of γ-irradiation and were given water containing Neomycin trisulfate (500 μg/ml, SIGMA N1876-25G) and Polymyxin (100U/ml SIGMA P1004-1MU)

### 4.2 Flow cytometry and cell sorting

Thymocytes were prepared by mincing the mouse thymus with a razor blade and pipetting the thymic fragment up and down. Before thymocyte staining, anti-mouse CD16/32 (Biolegend, Cat; 101302) was used as Fc Blocking. Thymocytes were stained with CD4-APCCy7 (Biolegend, #100526, diluted 1:400), CD8a-APC (Biolegend #100712, diluted 1:400), TCRb-PECy7 (Biolegend #109221, diluted 1:200), and PE-labeled CD1d-PBS-57 tetramer or CD1d-unloaded tetramer (diluted 1:400). Flow cytometric analysis and cell sorting were performed using BD FACSAria TM III (BD). Mouse CD1d-PBS-57 and control CD1d-unloaded tetramers were obtained from the Tetramer Core Facility, Emory University. Dead cells were excluded by 7-aminoactinomycin D staining. Data were analyzed using FlowJo TM 10 (Version 10.6.2). Flow cytometric analysis and cell sorting were performed using BD FACSAria III (BD).

### 4.3 RNA isolation and TCR Sequencing

Sorted thymocytes were suspended in Trizol (Invitrogen) to prepare total RNA according to the manufacturer’s protocol. NGS-based TCR repertoire analysis was performed by Repertoire Genesis Inc. (Osaka, Japan). Total RNA was converted to cDNA with Superscript III reverse transcriptase (Invitrogen). A BSL-18E primer containing polyT18 and a SphI site was used for cDNA synthesis. AftercDNA synthesis, double strand (ds)-cDNA was synthesized with Escherichia coli (E. coli) DNApolymerase I (Invitrogen), E. coli DNA ligase (Invitrogen), and RNase H (Invitrogen). ds-cDNAswere blunted with T4 DNA polymerase (Invitrogen). A P10EA/P20EA adaptor (Table S2) was ligated to the 5’ end of the ds-cDNA and then cut with SphI restriction enzyme. After removal of the adaptor and primer with MinElute Reaction Cleanup kit (Qiagen), PCR was performed with KAPA HiFi DNA polymerase (Kapa Biosystems, Woburn, MA) using TCRα (Tra) chain constant region-specific primer mCA1and P20EA. PCR conditions were as follows: 98°C (20 sec), 65°C (30 sec), and 72°C (1 min) for 20× cycles. The second PCR was performed with mCA2 and P20EA primers using the same PCR conditions. Amplicons were prepared by amplification of the second PCR products using P22EA-ST1-R and mCA-ST1-R.

After PCR amplification, index (barcode) sequences were added by amplification with Nextera XT Index kit v2 setA (Illumina, San Diego, CA). The indexed products were mixed in an equimolar concentration and quantified by a Qubit 2.0 Fluorometer (ThermoFisher Scientific). Sequencing was done with the MiSeq System (Illumina) using 300-bp paired-end reads (∼1.0×105 Tra and ∼1.8×105 Trb reads per condition).

### 4.4 Data analysis

All TCR-seq datasets were pre-processed by MiXCR (version 3.0.5; MiLaboratories Inc., San Francisco, CA) for TRAV and TRAJ genes assignment, and CDR3 extraction with parameters `-- species mmu --starting-material rna -receptor-type tra -5-end v-primers -3-end c-primers -adapters adapters-present`. Following analysis was performed in R (version 4.3.1). Unequal variances t-test for comparing means of groups was performed to the logarithm of the number of cells using ‘stats::t.test’ function in R.

### 4.5 Mathematical modeling

The mathematical model was simulated using ‘ParametricNDSolveValue’ function in Mathematica (version 12.0; Wolfram research, Champaign, IL). The parameters in the equation (1) were following; λ = 10, *n*_max_ = 1, *g*_1_ = 0.098, *g*_2_ = 0.002. We set the initial values *n*_1_ (0) = 0.1, *n*_2_ (0) = 0.01 for Fig. 5a and *n*_1_ (0) = 0.1, *n*_2_ (0) = 0.0001 for Fig. 5b.

## 5 Acknowledgments

We thank the members of our labs for the fruitful discussion. The authors acknowledge the use of ChatGPT-4 (OpenAI) and Grammarly (Grammarly, Inc.) to proofread the English language of the manuscript. The tools were applied exclusively for language refinement and did not contribute to the scientific content, interpretation, or conclusions. The authors reviewed and validated all AI-processed content to ensure accuracy and appropriateness.

## 6 Declarations

### 6.3 Author Contributions

**Kazumasa B. Kaneko**, Formal analysis, Investigation,Visualization, Writing - Original Draft Preparation; **Mio Hayama**, Data curation, Formal analysis; **Nanako Uchida**, Data curation, Formal analysis; **Takahisa Miyao**, Data curation, Formal analysis, Investigation, Validation; **Nobuko Akiyama**, Supervision, Investigation, Validation, Writing – review and editing; **Taishin Akiyama**, Investigation, Supervision, Validation, Writing – Original Draft Preparation, Writing - review & editing; **Tetsuya J. Kobayashi**, Formal analysis, Funding acquisition, Supervision, Validation, Writing - original draft preparation, Writing - review & editing;

### 6.4 Final Approval

All authors have reviewed the manuscript and agree to its submission.

### 6.5 Ethics Approval

The animal study was approved by Institutional Animal Care and Use Committee of RIKEN, Yokohama Branch. The study was conducted in accordance with the local legislation and institutional requirements.

### 6.6 Data Availability

The datasets and code for this study will be publicly available before publication.

