## Supplementary Figures for "TCR repertoire analysis quantifies effect of irradiation to homeostasis of iNKT cell development in the thymus of mice"

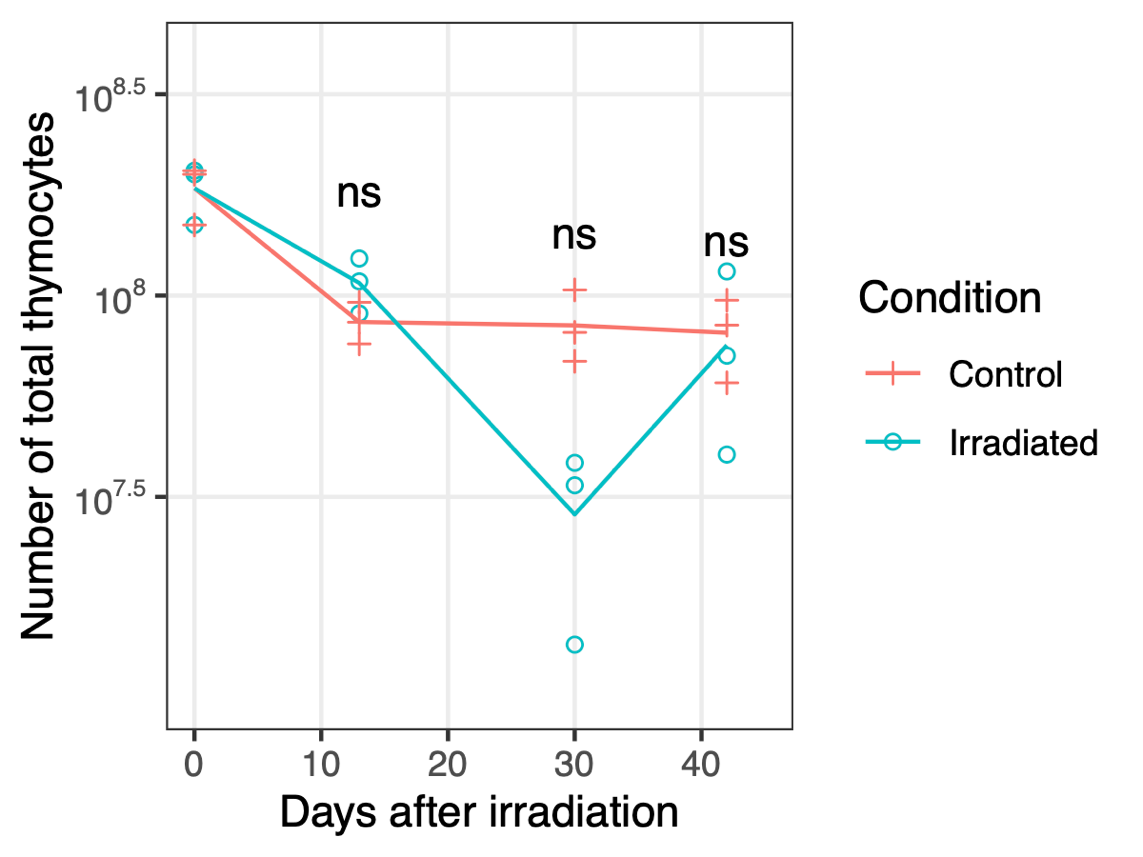

**Supplementary Figure 1.** Kinetics of total thymocytes after sub-lethal irradiation. 3 samples were analyzed at each time point before and after irradiation. Unequal variance t-test was conducted between irradiated and control groups at each time point. ns: p ≥ 0.05.
